# Ccr4 is a novel shuttle factor required for ubiquitin-dependent proteolysis by the 26S proteasome

**DOI:** 10.1101/2020.03.30.015370

**Authors:** Ganapathi Kandasamy, Ashis Kumar Pradhan, R Palanimurugan

## Abstract

Degradation of short-lived and abnormal proteins are essential for normal cellular homeostasis. In eukaryotes, such unstable cellular proteins are selectively degraded by the ubiquitin proteasome system (UPS). Furthermore, abnormalities in protein degradation by the UPS have been linked to several human diseases. Ccr4 protein is a known component of the Ccr4-Not complex, which has established roles in transcription, mRNA de-adenylation and RNA degradation etc. Excitingly in this study, we show that Ccr4 protein has a novel function as a shuttle factor that promotes ubiquitin-dependent degradation of short-lived proteins by the 26S proteasome. Using a substrate of the well-studied ubiquitin fusion degradation (UFD) pathway, we found that its UPS-mediated degradation was severely impaired upon deletion of *CCR4* in *Saccharomyces cerevisiae.* Additionally, we show that Ccr4 binds to cellular ubiquitin conjugates and the proteasome. In contrast to Ccr4, most other subunits of the Ccr4-Not complex proteins are dispensable for UFD substrate degradation. From our findings we conclude that Ccr4 functions in the UPS as a shuttle factor targeting ubiquitylated substrates for proteasomal degradation.

## Introduction

Normal cellular homeostasis requires the cells to regulate and/or degrade functional as well as non-functional proteins by proteolysis. In eukaryotes, the most important proteolytic system that is essential for selectively degrading such proteins is the ubiquitin proteasome system. Prior to degradation, the substrate proteins are marked with ubiquitin by a set of enzymes (E1, E2 and E3); then the ubiquitin-modified proteins are degraded by the 26S proteasome (Budenholzer et al., 2017; Dikic, 2017; Marques et al., 2009; Varshavsky, 2017). There are many diseases and abnormalities that are linked to defective protein degradation including parkinson’s, alzheimer diseases, cancer and muscular dystrophy etc. (Reinstein and Ciechanover, 2006; Rousseau and Bertolotti, 2018; Vinciguerra et al., 2010). The significance of UPS has been eminent after identification of many substrates of the proteasome including major regulatory factors such as cell cycle proteins (Cyclins) and signaling proteins (Rose and Mayor, 2018). Groundbreaking studies have lead to the characterization of specific protein degradation pathways in eukaryotes, such as the N-end rule (Bachmair et al., 1986; Dohmen et al., 1991; Ghislain et al., 1996; Madura et al., 1993) and ubiquitin fusion degradation pathways (Johnson et al., 1995), both of which depend on dedicated enzymes (E1, E2, E3 and DUBs) involved in ubiquitylation and de-ubiquitylation. Further studies on the same line resulted in the identification of proteins relating to the function of the proteasome (Ramos et al., 1998; Varshavsky, 2011). Several Ufd proteins, for example, Ufd1 to Ufd5 are required for the ubiquitin-dependent degradation of linear ubiquitin fusion proteins in the UFD pathway in eukaryotes. Specifically, Ufd4 is an ubiquitin ligase (E3) (Johnson et al., 1995), together with the ubiquitin-conjugating enzymes Ubc4 and Ubc5 (E2s) attaches ubiquitin to the UFD substrates. In the next step, Cdc48^Ufd1/Npl4/Ufd2^ complex binds to mono and/or di-ubiquitin modified UFD substrates, then the U-box containing protein Ufd2 elongates ubiquitin chain thereby optimizing the chain length and topology for efficient recognition and degradation by the proteasome (Richly et al., 2005). In an interesting study, Ubr1 (E3 for N-end rule pathway substrates) was found to form a complex with Ufd4 to ubiquitylate the DNA repair protein Mgt1 in yeast (Hwang et al., 2010). Ufd3 is a 80 KDa protein required for the degradation of UFD as well as the N-end rule substrates by the proteasome (Hwang et al., 2010). Although the molecular function of Ufd3 is still largely unknown, it is clear that it binds to Cdc48 and required for ubiquitin recycling as the free ubiquitin levels were found to be low upon *UFD3* deletion (Johnson et al., 1995). Ufd5, also known as Rpn4, is a transcription factor that is involved in regulating expression of UPS genes and controls proteasome levels. Therefore, a deletion of *UFD5/RPN4* results in stabilization of various substrates that are destined to be degraded by the proteasome (London et al., 2004; Xie and Varshavsky, 2001).

Aside from ubiquitin ligases (E3s) that functions as monomers or dimers, there are several E3s found to be multimeric proteins, such as anaphase-promoting complex (APC or cyclosome), SCF or Ccr4-Not complex (Nakayama and Nakayama, 2006; Page and Hieter, 1999; Peters, 1998; Vodermaier, 2004). The Ccr4-Not complex is a highly conserved protein complex present in all eukaryotes (Collart and Panasenko, 2017). Although, the composition of Ccr4-Not complexes vary slightly in different organisms, in yeast the Ccr4-Not complex is composed of at least nine subunits namely Ccr4, Caf1, Caf40, Caf130, Not1, Not2, Not3, Not4 and Not5 (Chen et al., 2001; Collart, 2003). The Ccr4-Not complex has multiple functions ranging from mRNA de-adenylation, RNA degradation, transcriptional regulation to translational control etc. (Inada and Makino, 2014)(Collart and Panasenko, 2017). Additionally, the Ccr4-Not complex is known to interact with several protein complexes including proteasome (Laribee et al., 2007) (Figure 7), ribosome and molecular chaperones (Preissler et al., 2015; Villanyi et al., 2014). Interestingly, the shared nuclease catalytic center in Ccr4-Not complex is composed of Ccr4, Caf1 and Not1 subunits (Basquin et al., 2012; Petit et al., 2012). The Not4 subunit contains a canonical RING motif, which is found in a class of ubiquitin ligases, and required for degradation of several proteasome substrates (Cooper et al., 2012; Gulshan et al., 2012; Mersman et al., 2009; Panasenko et al., 2006; Panasenko and Collart, 2012; Simonetti et al., 2017). However, the direct involvement of Not4 in ubiquitin modification of proteasome substrates has not been demonstrated so far. A recent study showed that Not4 can modify the Rpt5 subunit in the 19S regulatory particle (RP) of the 26S proteasome by monoubiquitylation *in vitro* (Fu et al., 2018). Ubiquitin modified Rpt5 is not subjected to protein degradation. Furthermore, in higher eukaryotes Not4 is not a part of the Ccr4-Not complex and there are a few additional subunits that are specifically found only in *Drosophila* and human Ccr4-Not complexes (Collart and Panasenko, 2017; Temme et al., 2010) suggesting that the precise molecular function of Not4 in protein degradation remains to be explored. Further investigations found that Not2, Not3 and Not5 subunits were required for 5’ de-capping of mRNA prior to its degradation (Alhusaini and Coller, 2016). Although the exact molecular functions of Caf40 and Caf130 are still unknown, these polypeptides are implicated in stabilizing the interactions in the Ccr4-Not complex, thereby contributing to its de-adenylation and ubiquitylation functions (Keskeny et al., 2019). Importantly, the Ccr4-Not complex also has been implicated in several diseases such as obesity, diabetes and liver steatosis (Morita et al., 2011). Additionally, the Ccr4-Not complex also has a role in energy metabolism (Morita et al., 2011). In an intriguing earlier observation, the deletion of *CCR4* and *CAF1* showed similar phenotypes when compared to the other *NOT* gene deletions found in Ccr4-Not complex (Halter et al., 2014). On the other hand, *NOT* gene deletions showed similar phenotypes when compared among the NOT genes suggesting that the Not proteins might also form a sub complex apart from being part of Ccr4-Not complex. Similarly, Ccr4 and Caf1 might also constitute a distinct complex that is independent of the Ccr4-Not complex (Winkler and Balacco, 2013). Ccr4 has earlier been shown to bind to the regulatory particle (RP) of the 26S proteasome indicting that Ccr4 might have role in protein degradation (Laribee et al., 2007) (Figure 7). Such role, however, has not been demonstrated until now. In this study, we investigated the role of the Ccr4-Not complex in degradation of ubiquitin-dependent as well as ubiquitin-independent substrates by the 26S proteasome. Our findings let us to conclude that Ccr4 is required for degradation of ubiquitin-dependent substrates. Upon deletion of *CCR4*, the UFD substrate degradation is severely impaired, and the *CCR4* mutant accumulates UFD substrate in the ubiquitin-modified form. From our results, we hypothesize that Ccr4 might function as a shuttle and/or targeting factor for ubiquitin-modified proteasome substrates.

## Materials and Methods

### Yeast media

Complete yeast peptone dextrose (YPD) and synthetic dextrose (SD) minimal media having 2% dextrose (glucose or galactose) with specific dropouts were prepared as described earlier (Ramos et al., 1998) (Palanimurugan et al., 2004) and used in this entire study.

### Strains and plasmids

BY4741 wild type yeast strain and its derivatives were used in this study are listed in the strain table attached in the supplemental information. The *TRP1* gene was deleted in BY4741 and *ccr4Δ* strains using a technique described earlier (Alani et al., 1987). In order to do that, pNKY1009 plasmid was digested with BglII and EcoRI to release a deletion cassette. The released fragment was used to transform respective yeast strains and Ura^+^ and Trp^-^ colonies were selected on appropriate media. In the following step, the URA3 gene was cured out by streaking out the selected colonies on SD medium containing (1g/l) 5-Fluoroorotic acid hydrate (Sigma) plates as described previously (Boeke et al., 1984). The genotypes of the resulting PYGA12 and PYGA14 strains are listed in the supplementary strain table 1A.

**Table 1A:** Strains

| Yeast strain | Genotype | Source |
| --- | --- | --- |
| BY4741-WT | *MATa his3-Δ1 leu2Δ0 met15-Δ0 ura3-Δ0* | Euroscarf |
| *ccr4Δ* | *MATa his3-Δ1 leu2Δ0 met15-Δ0 ura3-Δ0 ccr4::KanMX4* | Euroscarf |
| *ump1Δ* | *MATa his3-Δ1 leu2Δ0 met15-Δ0 ura3-Δ0 ump1::KanMX4* | Euroscarf |
| *not3Δ* | *MATa his3-Δ1 leu2Δ0 met15-Δ0 ura3-Δ0 not3::KanMX4* | Euroscarf |
| *not4Δ* | *MATa his3-Δ1 leu2Δ0 met15-Δ0 ura3-Δ0 not4::KanMX4* | Euroscarf |
| *not5Δ* | *MATa his3-Δ1 leu2Δ0 met15-Δ0 ura3-Δ0 not5::KanMX4* | Euroscarf |
| *caf30Δ* | *MATa his3-Δ1 leu2Δ0 met15-Δ0 ura3-Δ0 caf30::KanMX4* | Euroscarf |
| *caf140Δ* | *MATa his3-Δ1 leu2Δ0 met15-Δ0 ura3-Δ0 caf140::KanMX4* | Euroscarf |
| *not1-ts* | *MATa his3-Δ1 leu2Δ0 met15-Δ0 ura3-Δ0 not1-ts::KanMX4* | Euroscarf |
| *not2-ts* | *MATa his3-Δ1 leu2Δ0 met15-Δ0 ura3-Δ0 not2-ts::KanMX4* | Euroscarf |
| *ufd4Δ* | *MATa his3-Δ1 leu2Δ0 met15-Δ0 ura3-Δ0 ufd4::KanMX4* | Euroscarf |
| YAP16 | *MATa his3-Δ1 leu2Δ0 met15-Δ0 ura3-Δ0 trp1Δ KanMX4::P<sub>GAL1</sub>-PRE2* | This study |
| PYGA12 | *MATa his3-Δ1 leu2Δ0 met15-Δ0 ura3-Δ0 trp1Δ* | This study |
| PYGA14 | *MATa his3-Δ1 leu2Δ0 met15-Δ0 ura3-Δ0 trp1Δ ccr4::KanMX4* | This study |

**Table 1B:**
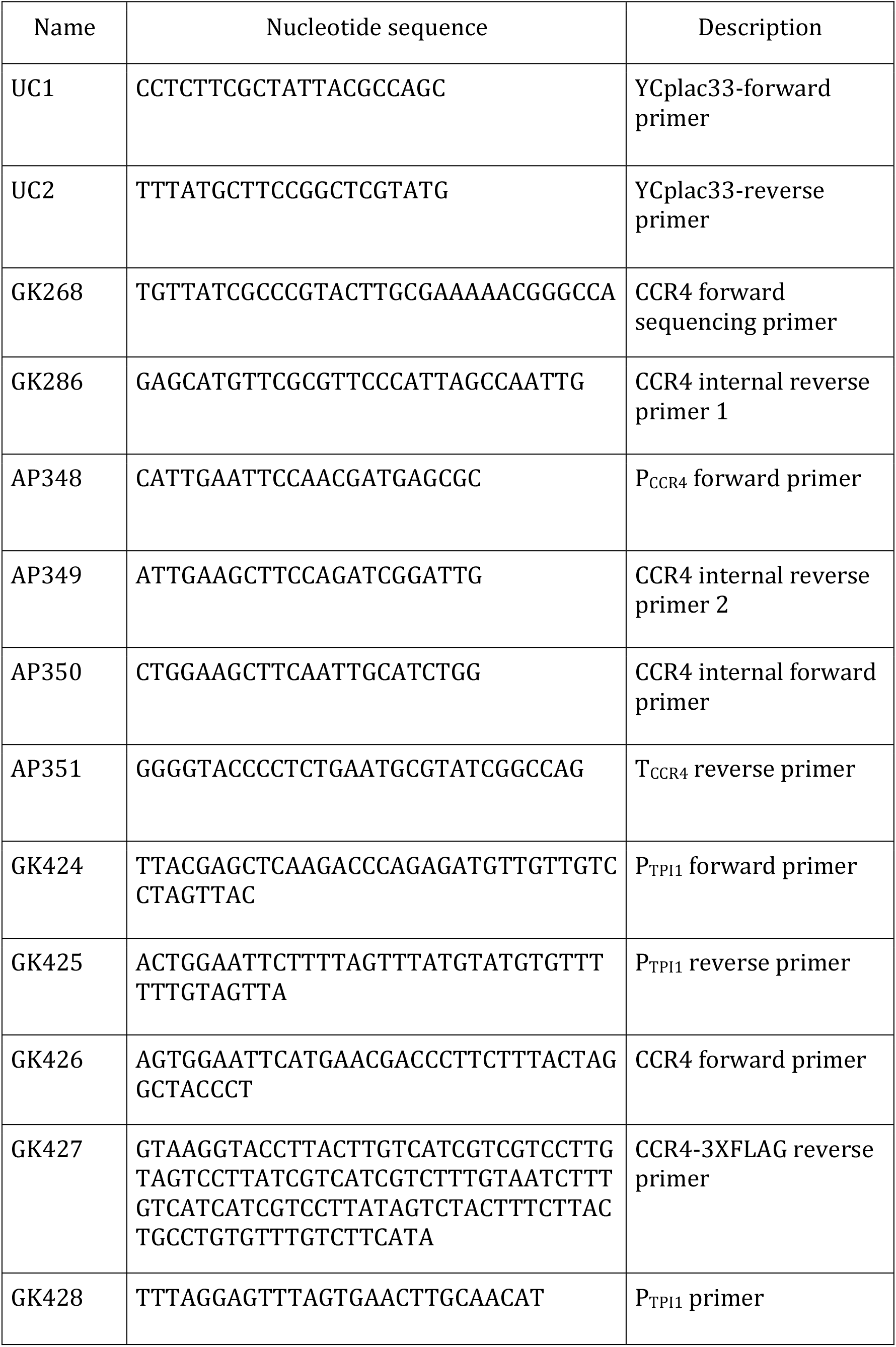
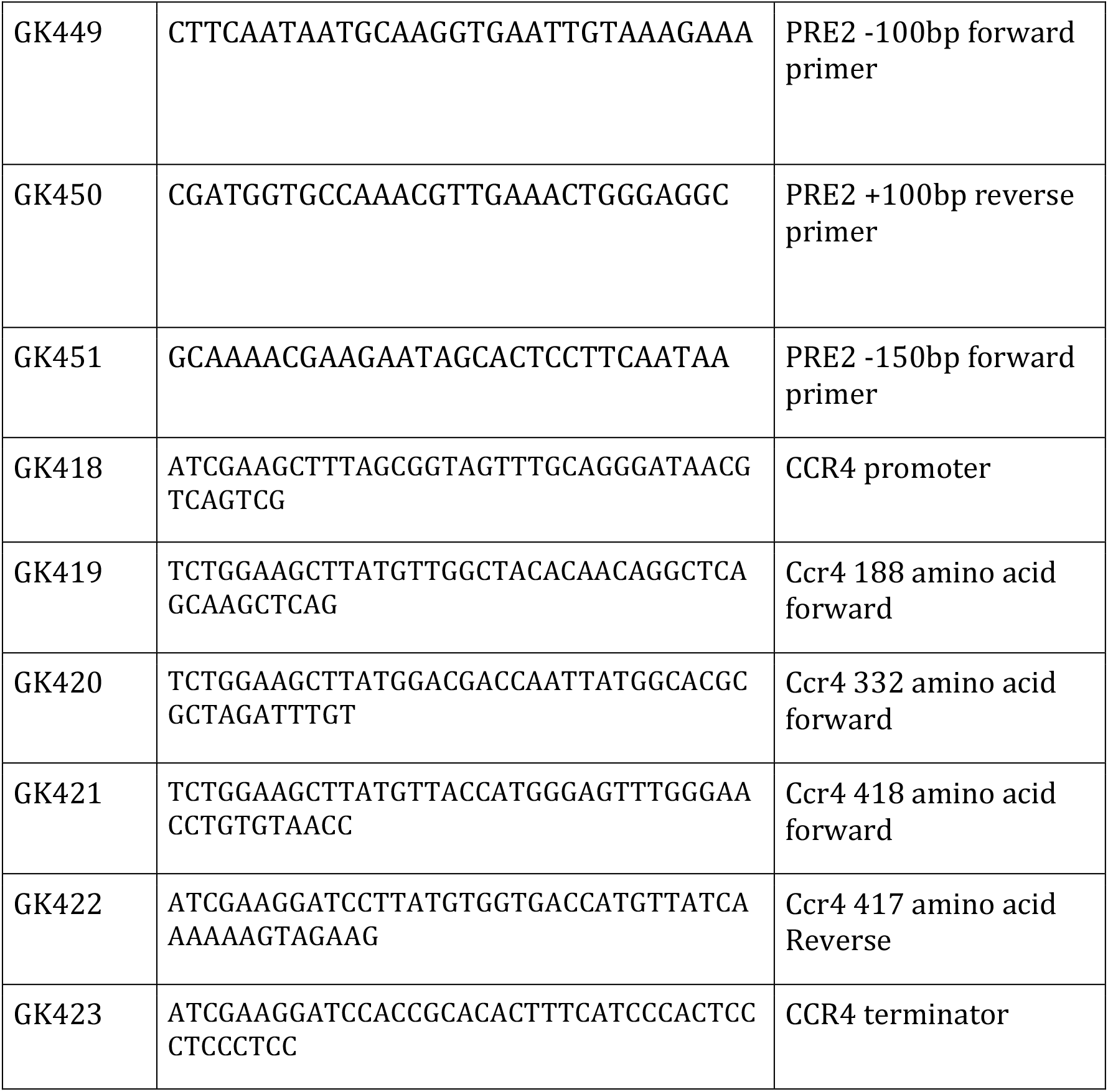
Primers

**Table 1C:**
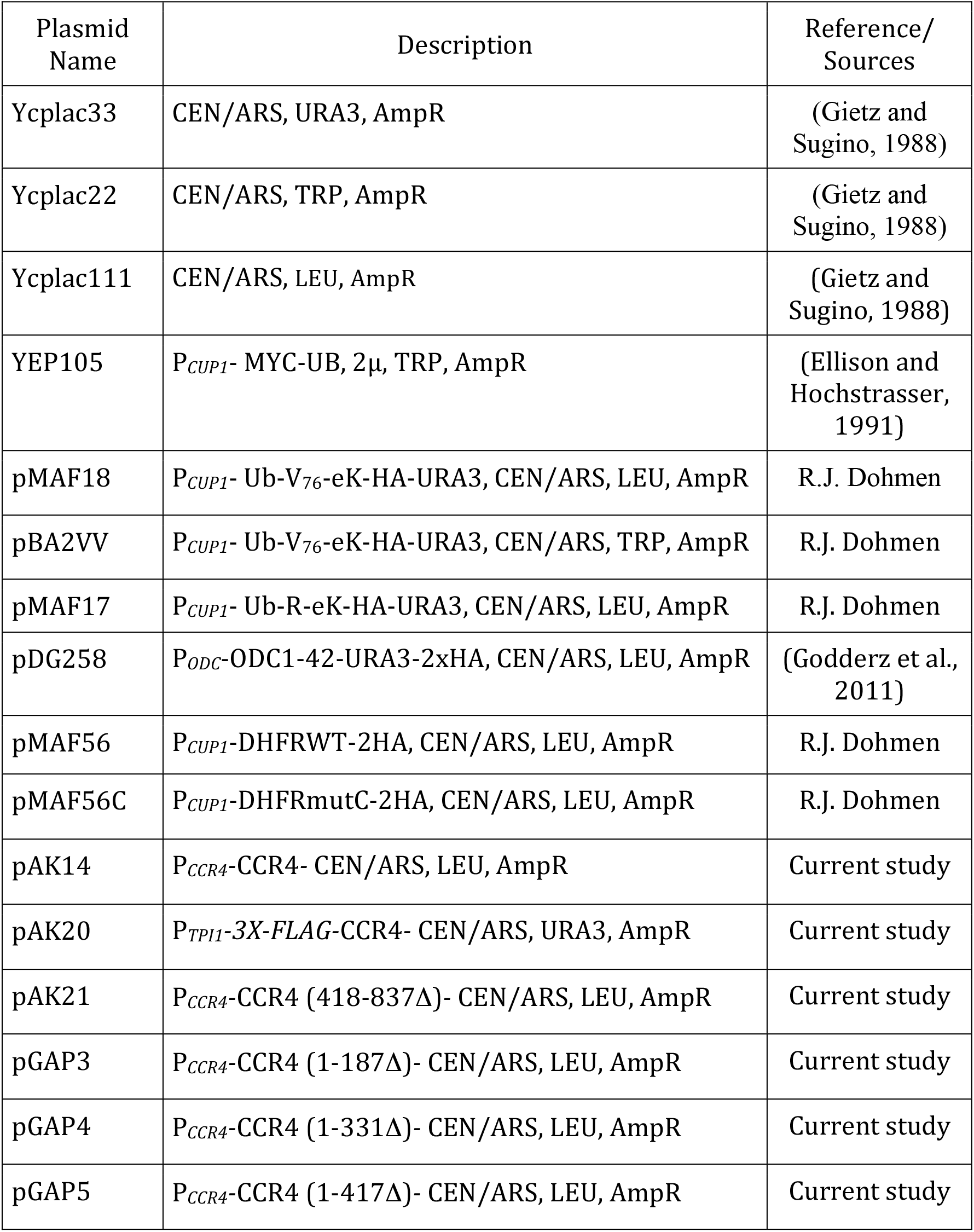
Plasmids

YAP16 strain was generated integrating a KanMX4::*P_GAL1_*-PRE2 cassette which was PCR amplified using the PRE2 gene flanking primers (GK449 and GK450) from genomic DNA isolated from YGA140 strain. PCR amplified DNA was used to transform PYGA12, and the transformants were selected on YPGal-G418 plate (300μg/ml). Plasmids pMAF17 (*P_CUP1_*-UB-R-eKa-HA-URA3), pMAF18 (*P_CUP1_*-UB-V^76^-eKa-HA-URA3), pMAF56C (*P_CUP1_*-DHFR^mutC^-2HA) and pMAF56 (*P_CUP1_*-DHFR^WT^-2HA) are derived from the *CEN/LEU2* vector pRS315 (Sikorski and Hieter, 1989) and pBA2VV is derived from *CEN/TRP1* vector pRS314, were kind gifts from R.J. Dohmen. pDG258 (*P_SPE1_*-ODS-URA3-2HA) is also derived from yeast *CEN/LEU2* vector pPM91 as described (Godderz et al., 2011; Kurian et al., 2011). pAK14 (*P_CCR4_*-CCR4-*T_CCR4_*) is derived from *CEN/LEU2* vector Ycplac111(Gietz and Sugino, 1988). This vector was digested with EcoRI and KpnI, then the *CCR4* fragment-1 containing promoter and a 5’ part of the *CCR4* ORF was generated using primers AP348 and AP349 and digested with EcoRI and HindIII, and a *CCR4* fragment-2 containing a 3’ part of *CCR4* ORF and terminator was generated using primers AP350 and AP351 and digested with HindIII and KpnI then used for ligation. For generating CCr4 deletion constructs *CEN/LEU2* vector Ycplac111 was used as a vector backbone. To construct Ccr4 1-187*4*, 331-837*4*, and 1-417*4* the insert 1 carrying the CCR4 promoter region was PCR amplified and double digested with EcoRI and HindIII. Next, the insert 2 either carrying nucleotides corresponding to Ccr4 188-837 (pGAP3), 332-837 (pGAP4) or 418-837 (pGAP5) amino acid was PCR amplified and double digested with Hind III and KpnI and finally ligated to Ycplac111 digested with EcoRI and KpnI restriction enzymes. For construction of Ccr4 418-837*Δ*, the insert 1 carrying the CCR4 promoter and nucleotides corresponding to 1-417 amino acids of CCR4 (pAK21) was PCR amplified and double digested with E.coRI and BamHI, and the insert 2 carrying the CCR4 terminator region was PCR amplified and digested with BamHI and KpnI and finally ligated to Ycplac111 digested with E.coRI and KpnI. pAK20 (*P_TPI1_*-CCR4-3XFLAG) is derived from the *CEN/URA3* vector pGAP1. The pGAP1 vector was digested with SacI and KpnI and ligated to equally digested *P_TPI1_*-CCR4-3XFLAG. The inserts were derived by PCR amplification using the primers listed in the supplementary table 1A. YEp105 (expressing *P_CUPI_*-MYC-UB) is a *2μ/TRP1* plasmid (Ellison and Hochstrasser, 1991). All the primers and plasmid details are listed in the supplementary table 1B & respectively.

### Yeast growth condition

Yeast strains were grown on YPD or SD agar plates at 25°C. For growth in YPD or SD liquid media, yeast cultures were incubated at 25°C with shaking at 180 rpm. For temperature-sensitive mutants, the growth conditions are described for the respective experiments in the results section.

### Western blotting analysis for protein detection and quantification

To assay the steady state levels of the substrate proteins, *S. cerevisiae* transformants were inoculated and grown overnight at 25°C in SD medium lacking leucine (SD-Leu) as described earlier. The primary cultures were then diluted in fresh SD-Leu medium (OD_600_ – 0.2), and the cells corresponding to 5.0 OD ware harvested at mid log phase (OD_600_ – 0.8 to 1.0). The cell pellets were washed once with water then mixed with 250 μl of ice cold 1.85N NaOH. After incubation for 10 min on ice, 250 μl of 50% TCA was added to each sample. Then cells were pelleted by centrifuging at 14,000 rpm for 10 min, the supernatant was discarded, the cells were re-suspended in 100 μl of 1M Tris (pH 7.5), and pelleted again as mentioned above. In the next step, the pellet was re-suspended in 1X Laemmli loading buffer (0.0625M Tris-HCl (pH 6.8), 2% SDS, 1% β-mercaptoethanol, 10% glycerol, 0.002% bromophenol blue) and lysed by boiling at 85°C for 5 min. After a brief centrifugation, the yeast cell extracts corresponding to 0.416 OD_600_ were loaded on to a gel, then SDS-PAGE and western blotting were done to detect the specific proteins as well as to quantify the corresponding bands as described earlier (Palanimurugan et al, 2004, Kurian et al, 2011). Anti-Ha (C29F4 from cell signaling technologies, 1:2000 dilution) was used for detecting ha-tagged proteins. Anti-Tpi1 (Lab source, 1:10000 dilution) and Anti-Pgk1 (Novex, 1:5000 dilution) were used to detect Tpi1 and Pgk1. Proteins were detected using either secondary anti-mouse or anti-rabbit IgG coupled to peroxidase (Millipore, 1:5000 dilution), western bright ECL substrate (Advansta) was used for developing the blots. The signals were captured using a Bio-Rad ChemiDoc™ MP Imaging system and quantified using the software Bio-Rad Image Lab 6.0.1.

### Cycloheximide chase analysis for assaying protein stability

Protein stability was assayed by cycloheximide chase analysis. Exponentially growing yeast cells were treated with 100 mg/L cycloheximide; then samples were collected at appropriate time points. Samples were then washed, lysed and analyzed on SDS-PAGE followed by immuno-blotting as described above. Signals were quantified and protein degradation rates were calculated as mentioned (Kurian et al., 2011; Palanimurugan et al., 2004).

### Immunoprecipitation of Ccr4-Flag

Yeast strain YAP16 (BY4741, KanMX4::*P¢ALi*-PRE2) was either transformed with empty plasmids (YCplac111 and YEplac112) or its isogenic derivates expressing MYC-UB (YEp105) and CCR4-3XFLAG (pAK20). Transformants were selected on synthetic galactose media plates and primary cultures were grown in galactose medium until unless mentioned. For shutdown of expression from *P_GAL1_*, primary cultures were diluted into fresh liquid medium containing 2% glucose and grown at 25°C for 12 hrs. For immunoprecipitation, cells corresponding to 40 OD_600_ were harvested and the pellets were frozen at −65°C. For lysis, pellets were thawn and re-suspended in 400 μl of lysis buffer (50mM HEPES pH7.5, 150mM NaCl, 5mM MgCl2, 40mM N-Ethylmaleimide, EDTA-free protease inhibitor cocktail (Roche) and 2mM PMSF (Sigma); then 300 μl of acid washed glass beads (425-600 μM from Sigma) was added to the cell suspension, then tubes were vortexed 3X1 min each with incubation on ice for 2 min in-between. After clearing cell debris, the lysate was mixed with Triton X-100 (1% final concentration) and centrifuged at 14,000 rpm for 10 min at 4°C. Equal amounts (Protein estimation by Bradford) of soluble fractions were incubated with 40 μl of pre-equilibrated anti-FLAG M2 affinity agarose gel (Sigma) in binding buffer (Lysis buffer + 1% Triton X-100). Binding was done at 4°C for 2 hrs; then beads were washed 3X5 min each with binding buffer. Ccr4-3XFLAG protein was specifically eluted using 70 μl elution buffer (binding buffer plus 35 μg of 3X FLAG peptide from sigma). Eluted materials were boiled and analyzed by SDS-PAGE. Anti-Flag (Sigma, 1:2000 dilution) or anti-Myc (9B11, Cell Signaling Technologies, 1:1000 dilution) was used for protein detection. Immunoprecipitation of Ccr4-3XFLAG was also done under non-native conditions. For this purpose, soluble cell lysates were prepared as described above and SDS (2%) as well as β-mercaptoethanol (1%) were added. The samples were then boiled at 95°C for 5 min. Next, the samples were diluted to reduce the SDS and β-mercaptoethanol to 0.05% and 0.025% respectively. Diluted samples were incubated with 60 μl of pre equilibrated antiFLAG M2 affinity agarose gel and washed prior to elution as described above. Ccr4-3XFLAG was eluted using 80 μl of 3X Flag peptide (40 μg in binding buffer).

### Co-immunoprecipitation of Ccr4-Flag and Rpt5

Yeast strain YAP16 carrying either empty vector plasmid YCplac111 or pAK20 plasmid expressing Ccr4-3XFLAG was grown till mid-log phase at 25°C in galactose containing medium. Cells corresponding to 100 OD_600_ were harvested and stored at −65°C until further use. Cell pellets were resuspended in 600 μl of lysis buffer and the soluble extracts were prepared as described in the previous section. The Ccr4-3XFLAG protein was bound to anti-FLAG M2 affinity agarose gel as mentioned earlier in the native co-immunoprecipitation protocol. After extensive washing the bound proteins are eluted and co-purification of Rpt5 with CCr4-FLAG protein was detected by using western blotting.

## Results

### Certain proteasome substrates accumulate upon deletion of *CCR4*

In *Saccharomyces cerevisiae,* the Ccr4-Not complex is composed of several subunits including Not and Caf polypeptides (see introduction). Distinct phenotypes were observed for strains with *NOT* gene deletions compared to one with *CAF* or *CCR4* gene deletions. Not4, in yeast a subunit of the Ccr4-Not complex, is a RING finger ubiquitin ligase, which has been shown to interact with yeast proteasomes. In contrast, Not4 protein is not a stable subunit of the human and *Drosophila* Ccr4-Not complexes. Interestingly, another study revealed the interactions of remaining Ccr4-Not complex proteins Ccr4, Not1 and Not4 and these proteins were found to interact with 19S regulatory particle of the 26S proteasome (Laribee et al., 2007) (Figure 7). The above findings let us to hypothesize that the Ccr4-Not complex or its components might have functions in protein degradation by the UPS apart from its earlier well-characterized roles in RNA metabolism and translational control. We have particularly focused on the Ccr4 subunit of the Ccr4-Not complex in this study. Ccr4 has not been found to have any role in protein turnover despite the above-mentioned observation of its interaction with 19S RP. In order to test our hypothesis, we have expressed several known well-characterized UPS substrates (R-Ura3, Ub-V^76^-Ura3, Dhfr^mutC^, ODS-Ura3) that are degraded via ubiquitin-dependent and independent mechanisms by the 26S proteasome (Figure 1) in yeast, and assayed their steady state levels in wild type, *ccr4Δ* and *ump1Δ* strains. The latter is impaired in proteasome function due to the absence of the assembly and maturation factor Ump1 and served as a control (Ramos et al. 1998). As expected, our steady state level analysis showed that the *ump1Δ* mutant, relative to the wild type, accumulated all tested proteasome substrates, consistent with a general proteasomal defect (Figure 2A). Surprisingly, the UFD pathway substrate Ub-V^76^-Ura3 as well as the N-end rule pathway substrate R-Ura3 accumulated in the *ccr4Δ* mutant. Importantly, we observed that ectopic expression of wild type CCR4 or of CCR4-FLAG restored the Ub-V^76^-Ura3 substrate degradation in *ccr4Δ* strain (Figure 2B), confirming that the increased levels were indeed due to the absence of Ccr4. In contrast to these two proteolytic substrates, the levels of two other test proteins were unaffected by the deletion of *CCR4.* Specifically, levels of Dhfr^mutC^, a folding-deficient mutant version of mouse dihydrofolate reductase is known to be degraded in chaperone-mediated and ubiquitin-dependent manner, were not increased in *ccr4Δ* mutant. Moreover, also the levels of ODS-Ura3, a modified version of the Ura3 protein carrying the N-terminal degradation signal of ornithine decarboxylase (ODC degradation signal/ODS), which is known to be degraded in a ubiquitin-independent manner (Gödderz et al. 2011), were also similar in wild type and *ccr4Δ* strain. We also tested if *ccr44* would alter the transcription from P_CUP1_ by using stable Dhfr as a control. Our result showered that expression from P_CUP1_ was not altered in *ccr44* strain (Supplemental Figure 2). Together, these findings indicated that Ccr4 is required for efficient degradation of some but not all proteasomal substrates.

**Figure 1:**
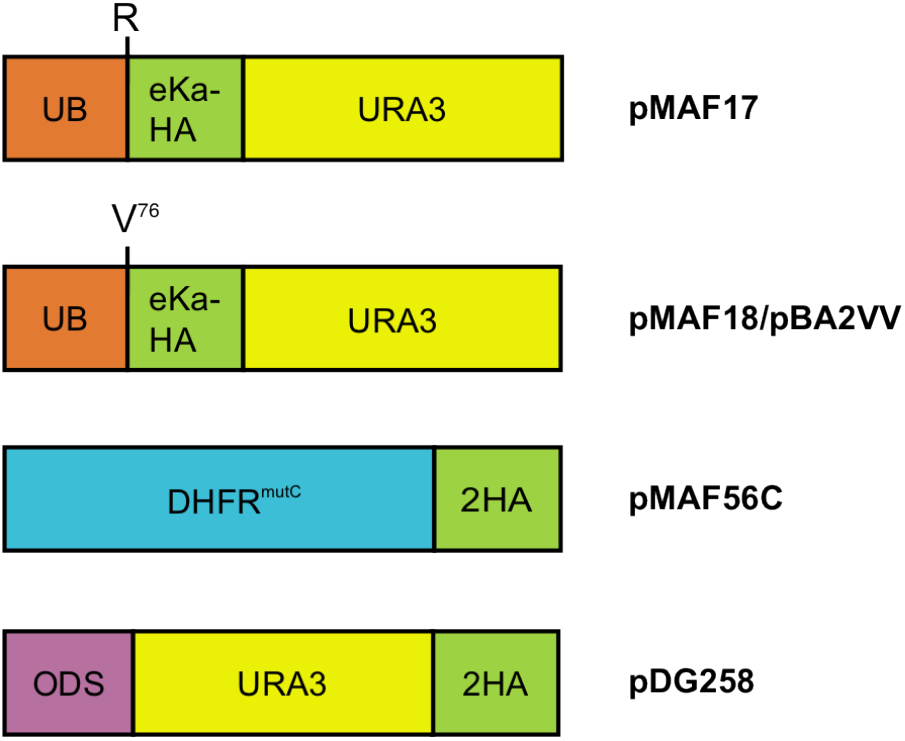
Illustration showing the various proteasome substrates. Cartoon depicting the salient features of proteasome substrates used in this study that are degraded by ubiquitin-dependent protein degradation pathway namely the N-end rule (pMAF17), ubiquitin fusion degradation (UFD) (pMAF18), quality control pathway protein mouse Dhfr^mutC^ (pMAF56C) and the ubiquitin-independent proteasome substrate ODS-Ura3 (pDG258).

**Figure 2:**
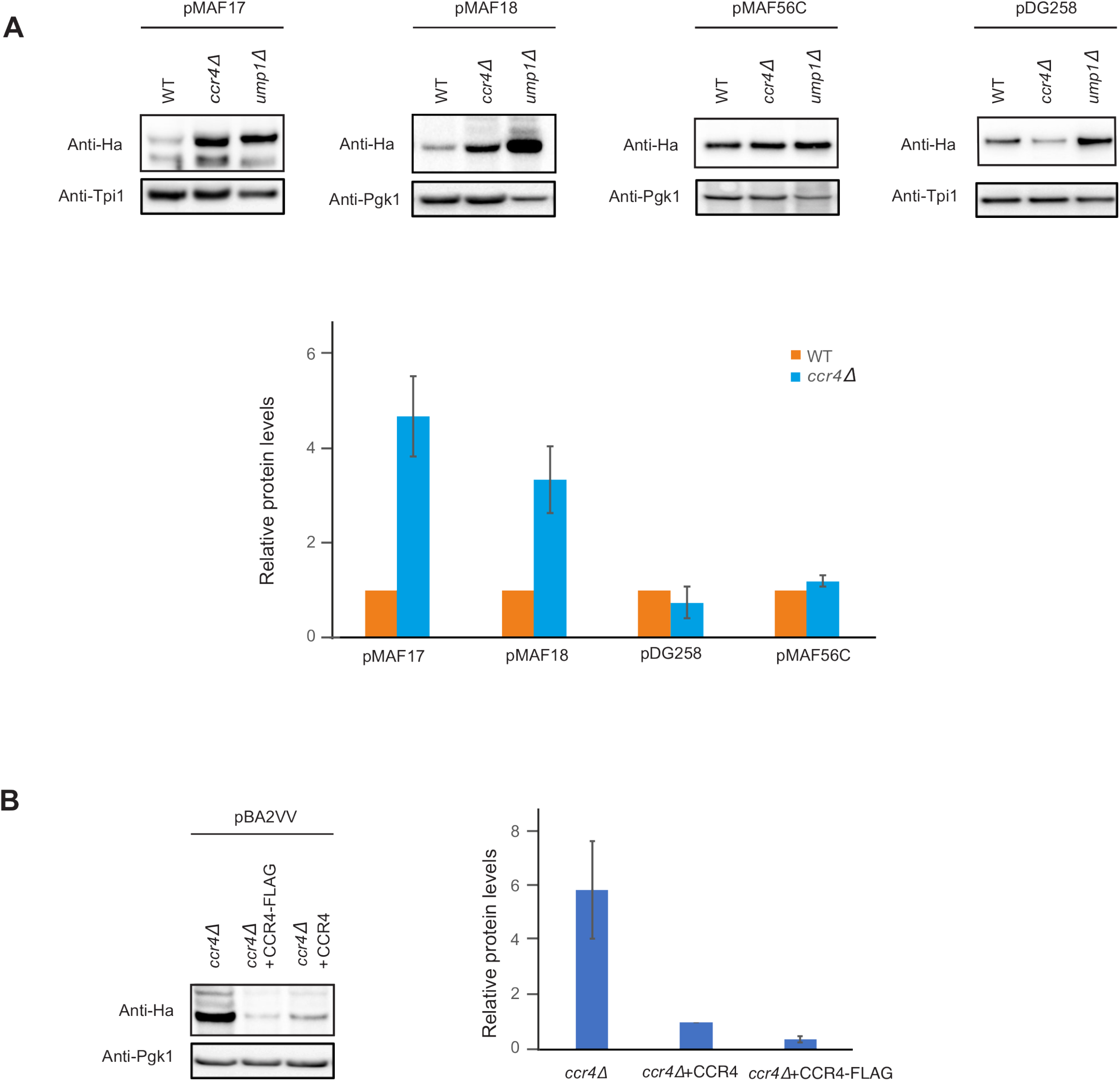
Steady state level analysis of proteasome substrates in Ccr4-Not complex subunit mutants. (A). Western blot analysis of various substrate levels in WT, *ccr4Δ* and *ump1Δ* strains. WT and mutant transformants having pMAF17, pMAF18, pMAF56C and pDG258 were grown in the SD-leu and with 100 μM CuSO_4_ till the cultures reached mid-log phase at 25°C. Cells were harvested and extract corresponding to 0.416 OD_600_ was analyzed by western blotting. Substrate proteins were detected using anti-Ha (upper panels). Tpi1 and Pgk1 were used as a protein loading control (lower panels). Note that in *ump1Δ* lanes the total protein loaded was less than WT and *ccr4Δ.* Quantification of signals from blots shown in panel above. Error bars denote s.d.; n≥3. (B). The *ccr4Δ* (PYGA14) strain carrying pBA2VV along with CCR4 (pAK14) and CCR4-3XFLAG (pAK22) gene variants were grown in the SD-leu-trp with 100 μM CuSO_4_ till the cultures reached mid-log phase at 25°C. Cells were harvested and extract corresponding to 0.416 OD_600_ was analyzed by western blotting. Substrate proteins were detected using anti-Ha (upper panels). Pgk1 was used as a protein loading control (lower panels). Quantification of signals from blots shown in panel B. Error bars denote s.d.; n≥3.

Considering the accumulation of certain proteasomal substrates in the *ccr4Δ* strain, we wanted to address the involvement of the remaining Ccr4-Not complex subunits in the substrate degradation. To this end, we have assayed the Ub-V^76^-Ura3 levels in *caf40Δ, caf130Δ, notl-ts, not2-ts, not3Δ and not5Δ* mutant strains. Our results show that Ub-V^76^-Ura3 levels did not change significantly compared to wild type levels in any of these mutants (Figure 3A). In addition, we monitored the growth of wild type, *ccr4Δ, ump1Δ*, and *ufd4Δ* strains expressing Ub-V^76^-Ura3 protein in medium lacking uracil. All these strains carry a mutation in the chromosomal *URA3* locus and require either sufficient amount of plasmid-encoded Ura3 protein or uracil in the media for normal growth. Only upon an impaired degradation of the above-mentioned Ura3-based test proteins, the strains should be able to grow on medium without uracil. Our assay revealed that expression of Ub-V^76^-Ura3 promoted growth of *ccr4Δ, ump1Δ* and *ufd4Δ* cells (Ufd4 is the E3 for UFD substrates)(Johnson et al., 1995) (Figure 3B) suggesting that Ub-V^76^-Ura3 accumulates upon *CCR4* deletion consistent with the observed increased steady state levels shown in figure 2.

**Figure 3:**
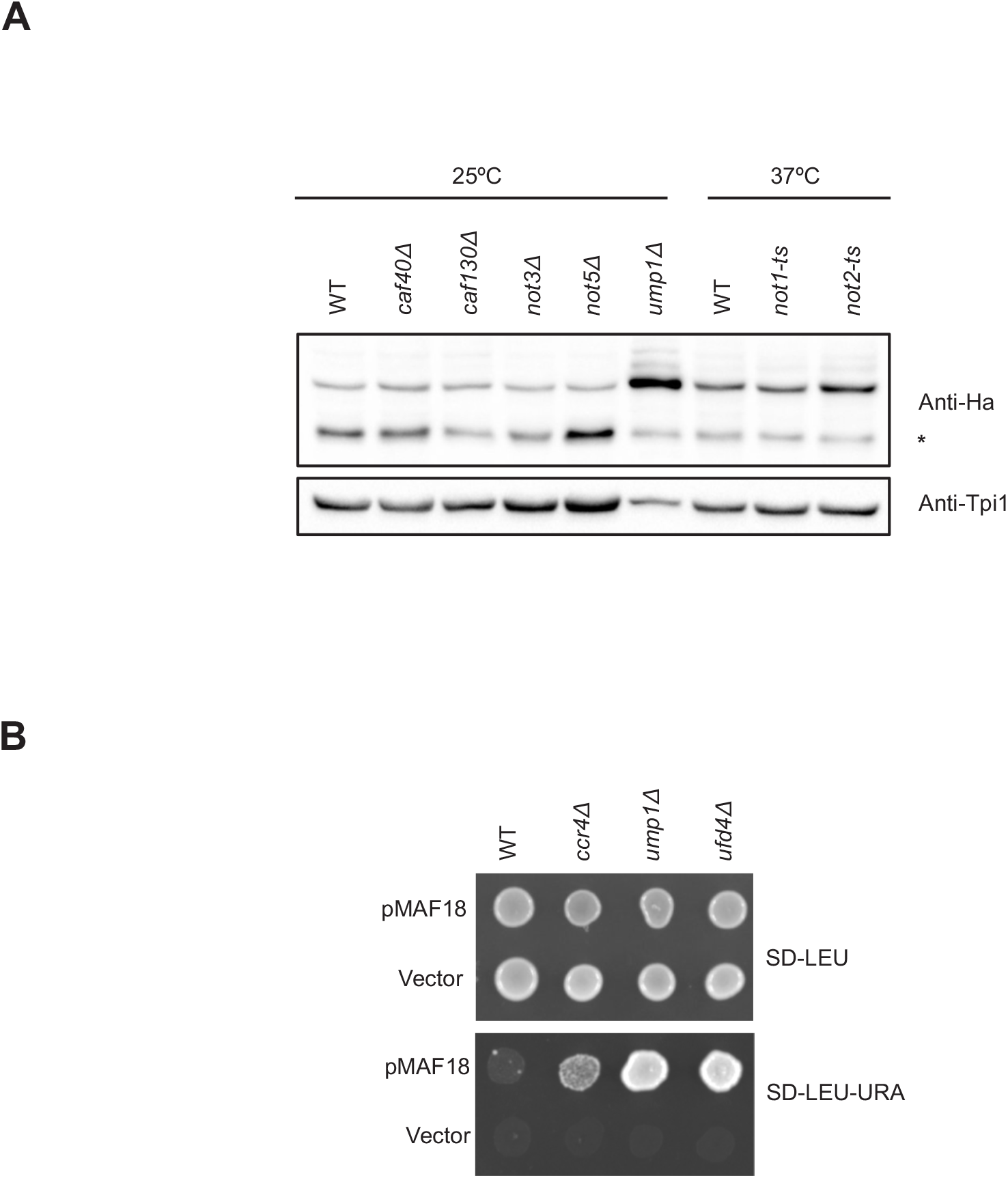
Steady state level analysis of UFD substrate in various Ccr4-Not subunit mutants. (A). Western blot analysis of UFD substrate (Ub-V^76^-Ura3) levels in various Ccr4-Not complex subunit deletion strains and temperature-sensitive mutants (*ts*). UFD substrate cassette was expressed from *P_CUP1_* by adding 100 μM CuSO_4_ to growth medium. Stable deletion strains were grown and harvested at 25°C and *ts* mutants *(notl-ts* and *not2-ts,* its isogenic WT strain) were initially grown at 25°C and later shifted to 37°C for 1 hr before harvesting. Upper panel shows anti-Ha signals and the lower panel is showing loading control Tpi1 signals. Asterisk denotes degradation product of UFD substrate. (B). Growth assay showing the accumulation of Ura3 based substrate proteins. WT, *ccr4Δ, ump1Δ,* and *ufd4Δ* having empty plasmid and pMAF18 were grown to mid-log phase. Then cells were spotted (3 μl from 10 OD_600_/ml suspension) on selection plates as indicated. Plates were incubated at 25°C for 4 days and photographed.

### UFD substrate turnover is impaired in the *CCR4* mutant

The observed increased steady state levels of Ub-V^76^-Ura3 in *ccr4Δ* mutant could be due to the following reasons. Firstly, the rate of synthesis of Ub-V^76^-Ura3 could have been altered at the transcription and/or translation steps resulting in increased protein levels. Alternatively, the stability of Ub-V^76^-Ura3 could have increased due to *CCR4* deletion. In order to identify the cause, we performed cycloheximide chase analyses to measure the Ub-V^76^-Ura3 turnover rates in wild type, *ccr4Δ* and *ump1Δ* strains. Surprisingly, we found that Ub-V^76^-Ura3 degradation was blocked upon deletion of *CCR4,* while the protein degradation was normal in the wild type strain (Figure 4A, B). The control *ump1Δ* strain showed even stronger stabilization of Ub-V^76^-Ura3. These observations suggest that Ccr4 is a novel factor essential for degradation of Ub-V^76^-Ura3. Similarly, we have performed cycloheximide chase analysis for ODS-Ura3 and Dhfr^mutC^. The results show that ODS-Ura3 as well as Dhfr^mutC^ were degraded normally in *ccr4Δ* strain (Supplemental Figure 1) suggesting that among the tested substrates Ccr4 is not required specifically for the degradation of ODS-Ura3 as well as Dhfr^mutC^.

**Figure 4:**
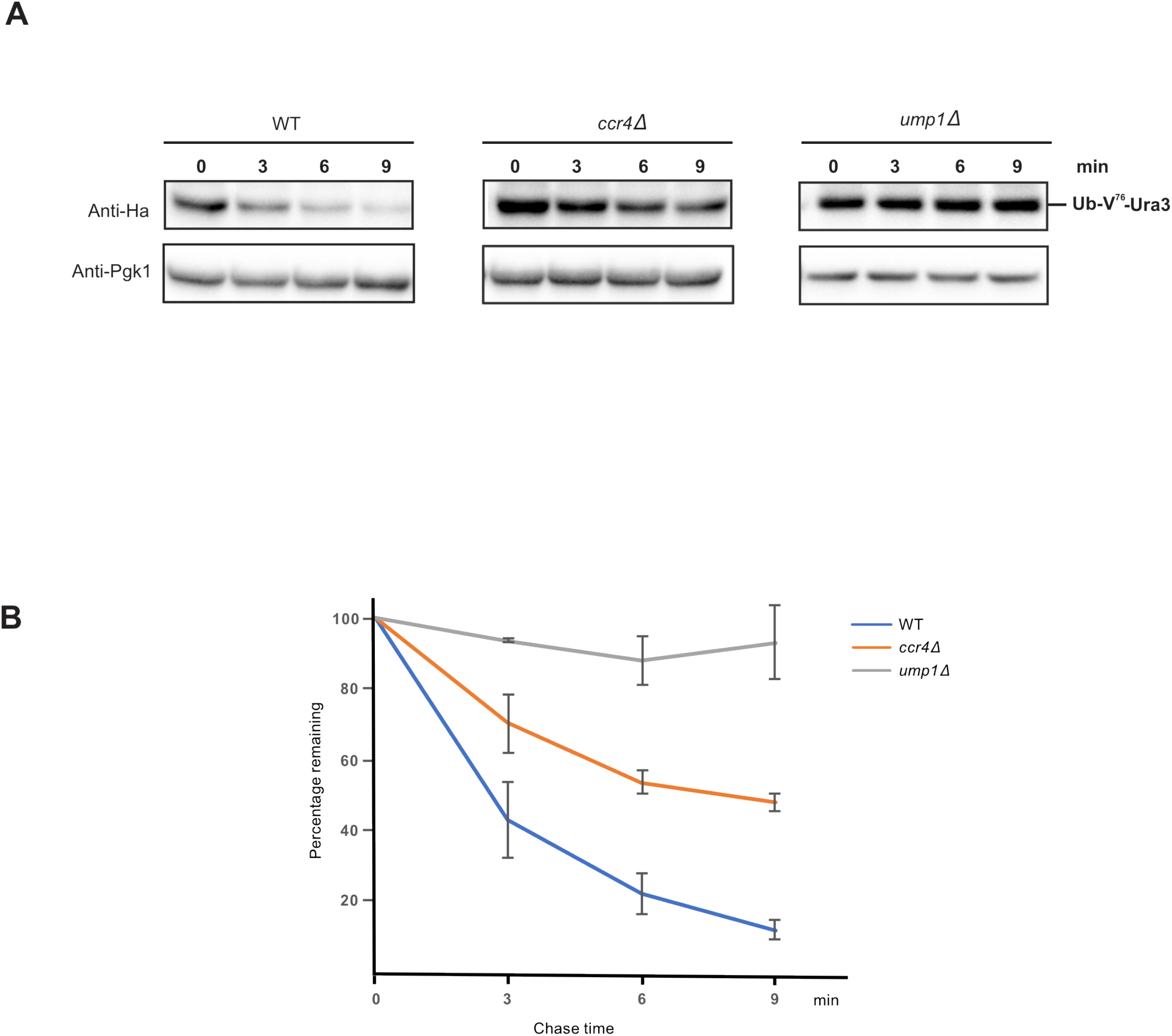
Cycloheximide chase assay for UFD substrate degradation. (A). WT, *ccr4Δ* and *ump1Δ* strains were transformed with pMAF18 expressing UFD cassette and the cells were grown in SD-leu with 100 μM CuSO_4_ till mid-log phase at 25°C. In the next step cycloheximide chase was done as mentioned in methods section and the samples were analyzed by western blotting. Anti-Ha was used for detecting the UFD substrate (Ub-V_76_-Ura3) (upper panels). Pgk1 served as a loading control (lower panels). (B). Quantification of the signals shown in panel. Error bars denote s.d.; n≥3.

### Ccr4 binds to cellular ubiquitin conjugates

Our finding that Ccr4 is required for efficient Ub-V^76^-Ura3 turnover raises an interesting question. How does Ccr4 promote UFD substrate degradation? To answer this question, we have considered several possible steps in UFD substrate degradation, in which Ccr4 might play a vital role. Broadly, all these steps could be classified in to two parts. A first part is substrate recognition and ubiquitylation and second part is the post-ubiquitylation substrate targeting and degradation. For Ub-V^76^-Ura3 the following enzymes are involved in ubiquitylation namely Uba1 (E1), Ubc4 and Ubc5 (E2s), Ufd4 (E3) and Cdc48^Npl4-Ufd1-Ufd2^ (processivity factor) for ubiquitin chain elongation. For targeting ubiquitin-modified substrates, there are several shuttle factors (such as UBA-UBL domain containing proteins Rad23, Dsk2 and Ddi1) that have been identified so far. Shuttle factors on the one hand binds to the ubiquitin chain of the modified substrates via their UBA domains, and on the other hand bind to the proteasome via their UBL domains thereby promoting substrate recognition and subsequent degradation by the 26S proteasome. In order to address the above possibilities, we analyzed the ubiquitylation status of Ub-V^76^-Ura3 in wild type, *ccr4Δ* and *ump1Δ* strains. Remarkably, our experiment showed that higher molecular weight ubiquitin-modified forms of Ub-V^76^-Ura3 accumulated upon deletion of *ccr4* similar to what was observed when *ump1* was deleted (Figure 5A) suggesting that Ccr4 is required for Ub-V^76^-Ura3 degradation at the post ubiquitylation step. Based on our result, we hypothesized that Ccr4 could be a novel shuttle factor for ubiquitin-modified substrates similar to those described above. To support our hypothesis, we tested whether Ccr4 interacts with 26S proteasome, indeed our coimmunoprecipitation experiments showed that the 19S regulatory particle subunit Rpt5 co-purified with Ccr4 protein (Figure 7), as reported earlier (Laribee et al., 2007). Next, we investigated if Ccr4 could also bind to cellular ubiquitin conjugates by co-immunoprecipitation assay. Excitingly, our results showed that ubiquitin conjugates co-immunoprecipitated along with Ccr4 under native conditions (Figure 5B). Upon denaturation of cellular ubiquitin conjugates, Ccr4 was unable to bind to ubiquitin conjugates (Figure 6). In conclusion, our results identified that the shuttling of ubiquitin-modified proteins to the 26S proteasome as a novel molecular function of Ccr4.

**Figure 5:**
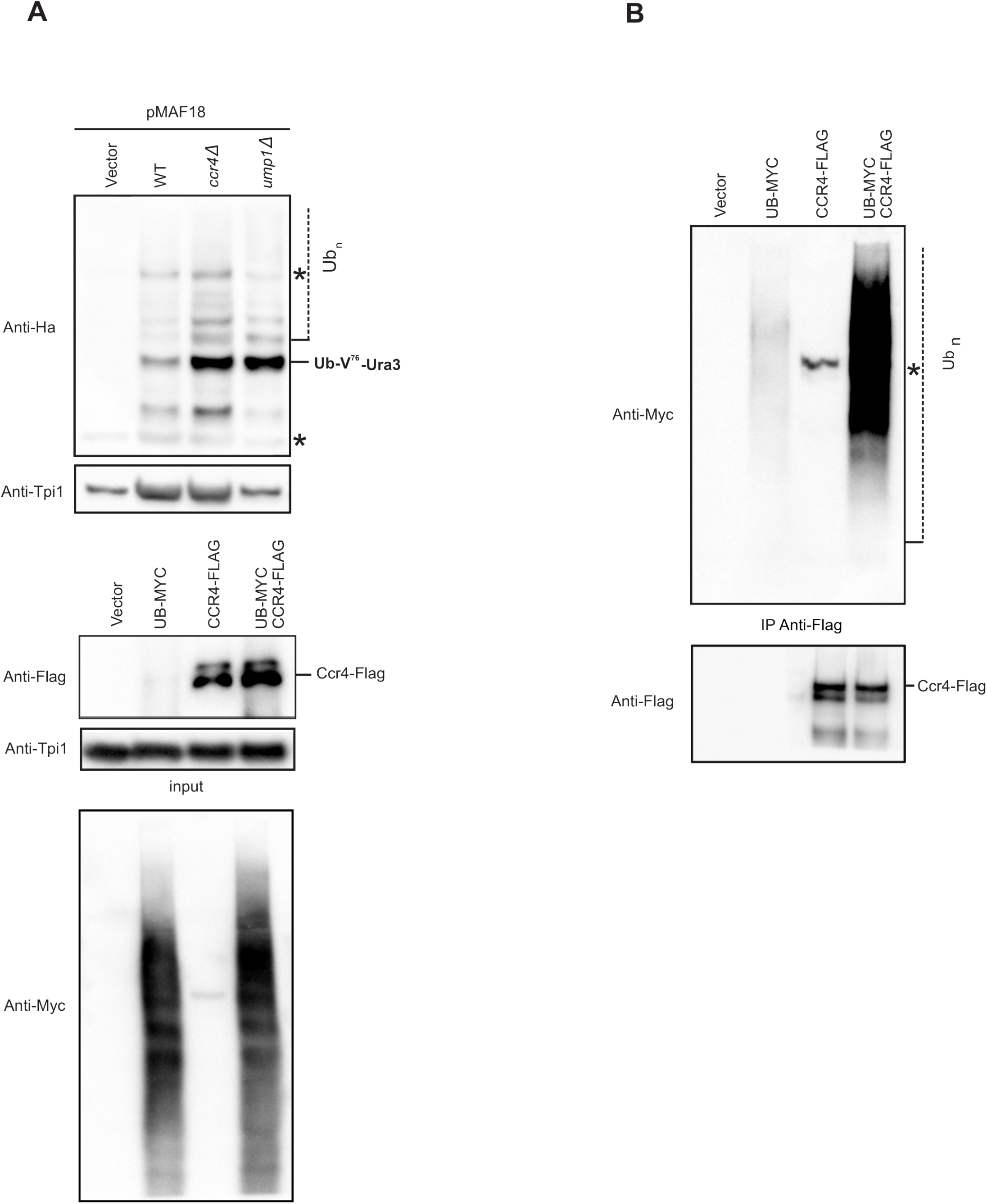
*CCR4* deletion accumulates ubiquitin modified UFD substrates and Ccr4 binds to cellular ubiquitin conjugates. (A). Western blot showing the higher molecular weight ubiquitin conjugated UFD substrate in WT, *ccr4Δ* and *ump1Δ*. Cells were grown at 25°C and samples were prepared as mentioned in methods section and loaded on low percent (8 %) SDS-PAGE gel to resolve the higher molecular weight forms. Anti-Ha was used for detecting UFD substrate protein (upper panel) and anti-Tpi1 was used for detecting the loading control (lower panel). Asterisk indicates nonspecific band. (B). Western blot showing association of higher molecular weight cellular ubiquitin conjugates with Ccr4-3XFLAG protein in native co-immuno precipitation experiment. YAP16 strain carrying either vector alone and/or plasmids expressing MYC-UBI, CCR4-3XFLAG was grown and Ccr4-3XFLAG protein was immunoprecipitated as described in methods section. Input levels (left panel) and eluted IP materials (right panel) were analyzed by western blots using anti-Myc and anti-Flag antibodies. Anti-Tpi1 was used to detect the loading control.

**Figure 6:**
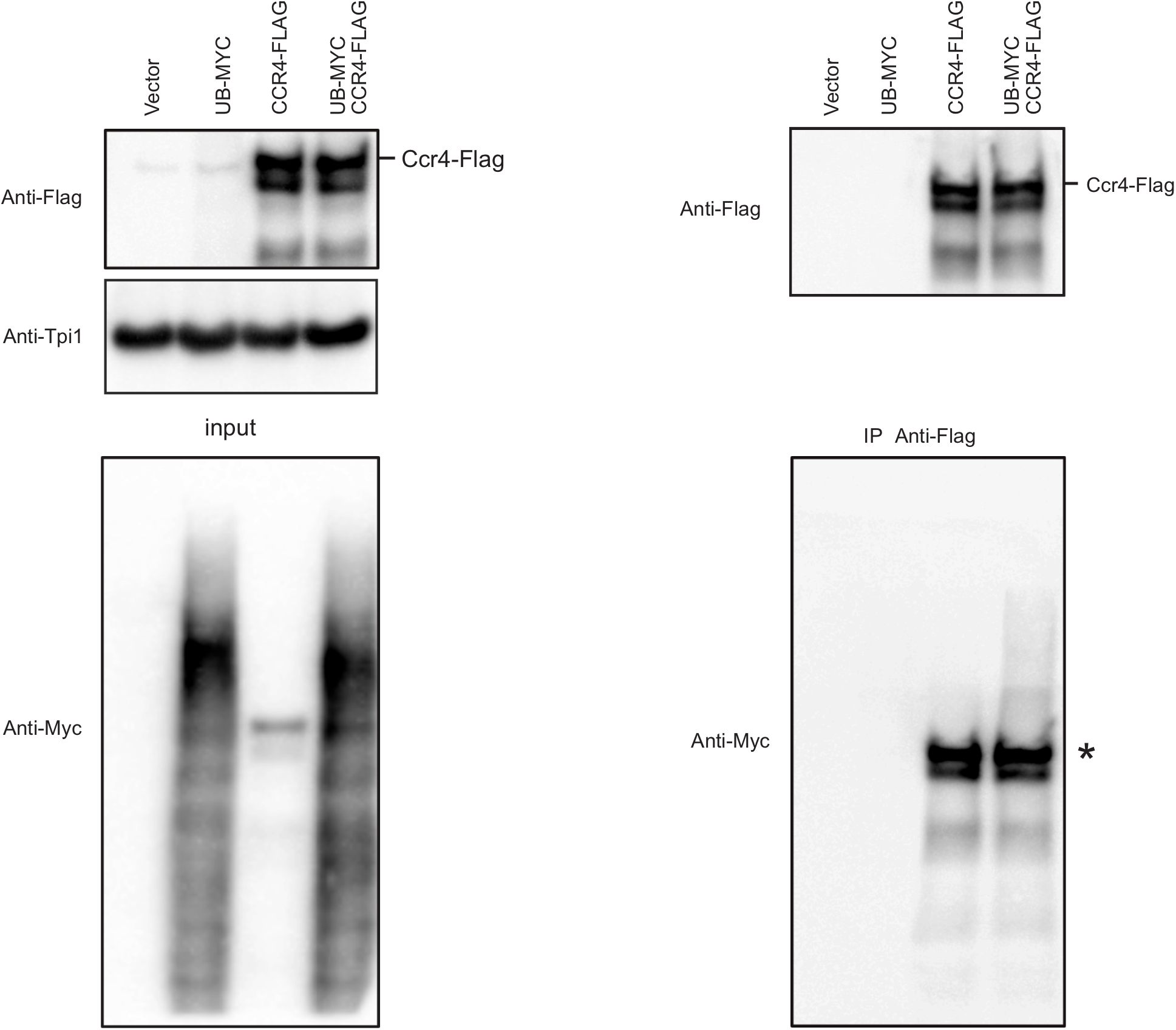
Ccr4 does not bind to cellular ubiquitin conjugates under denaturation conditions. IP Experiments were done as in Figure 5B except the lysates were subjected to denaturation by addition of 2% SDS, 1% β-mercaptoethanol and heating at 95°C for 5 min. Ccr4-3XFLAG protein was immunoprecipitated as described in methods section. Input levels (left panel) and IP eluate (right panel) were analyzed by western blots using anti-Myc and anti-Flag antibodies. Tpi1 was used as loading control.

### LRR and Nuclease domains of Ccr4 are indispensable to target UFD substrate for proteolysis

Ccr4 is an 837 amino acid residue long protein, it contains three major domains (Basquin et al., 2012). The residues 1 to 332 is rich in Q/N amino acids and it is known to form an unstructured region. Next, 333 till 470 region has five leucine rich repeats and constitutes the LRR domain of Ccr4 protein. From residues 471 to 837 is the c-terminal nuclease domain and executes mRNA deadenylation activity (Basquin et al., 2012). In order to find what region of Ccr4 protein contributes to ubiquitin shuttle activity, we generated several Ccr4 deletion constructs (Figure 8 A) and studied their effect on UFD substrate degradation. Our results revealed that the constructs expressing both the LRR and nuclease domain (188-837 and 332-837) complements the defective protein degradation phenotype of *ccr4Δ* mutant (Figure 8B&C). Intriguingly, the constructs expressing either the entire N-terminal/C-terminal region along with part of LRR (1-417 and 418-837 respectively) showed severe impairment in targeting UFD substrate for proteolysis (Figure 8B&C). The same observation was further confirmed by growth assay analysis using pBA2VV plasmid producing Ub-V^76^-Ura3 protein (Figure 8D). Altogether, these findings implicated that both LRR and nuclease domains of Ccr4 is essential for its function to target ubiquitin-modified protein for degradation.

## Discussion

Degradation of unstable proteins in eukaryotes are mainly carried out by the ubiquitin proteasome system and autophagy, which together result in recycling of amino acids and regulation of cellular homeostasis (Budenholzer et al., 2017; Dikic, 2017; Marques et al., 2009; Varshavsky, 2017). In this context, our study revealed that the Ccr4 protein is a novel shuttle factor in the UPS-mediated protein degradation. Ccr4 is an exonuclease known to be an active enzyme in the Ccr4-Not complex and has also been shown to physically interact with proteasome, the ribosome and the ribosome-bound chaperone NAC (nascent peptide associated complex) (Laribee et al., 2007)(Preissler et al., 2015; Villanyi et al., 2014). The biological significance of Ccr4 interaction with these protein complexes however is not established. In a recent study, deletion of *CCR4* was shown to result in higher amounts of ubiquitin-modified nascent proteins that are bound to yeast ribosomes (Duttler et al., 2013). Such an increase could be due to several reasons. Firstly, the degradation of ubiquitin-modified nascent proteins could be impaired upon *CCR4* deletion or the rate of ubiquitin deconjugation might be reduced. Further, there could have been increased rate of ubiquitin conjugation to nascent proteins associated with ribosomes in the absence of Ccr4 might be the cause of the observed increased levels of so modified polypeptides. However, there were no further studies to define the molecular function of Ccr4 among the above mentioned points (Duttler et al., 2013). In the present work we have used several well-characterized substrates of the proteasome that are degraded with ubiquitylation (R-Ura3, Ub-V^76^-Ura3, Dhfr^mutC^) or without ubiquitylation (ODS-Ura3). Our results showed that the UFD pathway substrate Ub-V^76^-Ura3 and the N-end rule pathway substrate R-Ura3 specifically accumulated in *CCR4* deletion strain, while the remaining tested substrates levels were similar to those in wild type (Figure 2). These observations led us to postulate that Ccr4 might be involved in UFD pathway and the same hypothesis was validated by cycloheximide experiments showing that the UFD protein stability was altered in *CCR4* mutant strain (Figure 4). While the remaining substrates listed above were degraded normally in *CCR4* mutant strain (Supplemental Figure 1), the UFD pathway substrate was stabilized upon the deletion of *CCR4.* From these results we conclude that the Ccr4 protein is a novel component of the UFD pathway and it is essential for Ub-V^76^-Ura3 degradation by the 26S proteasome. Interestingly, the other Ccr4-Not complex subunit deletion strains did not show any significant variations in Ub-V^76^-Ura3 levels (Figure 3A) suggesting that either Ccr4 works alone or in a subcomplex with other proteins to mediate UFD substrate degradation. This conclusion was re-validated with growth assays reporting on the intracellular levels of Ura3-based reporter proteins (Figure 3B). After discovering that the Ccr4 is required for UFD substrate degradation we wanted to understand at which step in the UFD pathway that Ccr4 promotes Ub-V^76^-Ura3 substrate degradation. On the one hand Ccr4 could affect the ubiquitylation of UFD substrate, on the other hand Ccr4 could affect the substrate targeting to the proteasome. In addition, Ccr4 could also affect proteasome activity. The observation that the degradation of a folding deficient quality control substrate mouse Dhfr^mutC^ is normal in *CCR4* deletion strain suggested that the substrate ubiquitylation is not affected in the absence of Ccr4. Additionally, the normal degradation of ODS-Ura3 by the proteasome is also suggesting that Ccr4 does not alter proteasome activity. Therefore the most likely step in which the Ccr4 could play a role appears to be the targeting of ubiquitin modified substrate to the proteasome. Our steady state blot analysis revealed that Ub-V^76^-Ura3 accumulates high molecular weight ubiquitin conjugates in the *ccr4Δ* mutant cells (Figure 5A). Therefore, we conclude that Ccr4 has a novel molecular function as a targeting factor for ubiquitin-modified proteins to the proteasome. As listed in the introduction there are several proteins known to be involved in targeting ubiquitin-modified substrates to the proteasome. Rpn1, Rpn10 and Rpn13 subunits of the 19S caps of the proteasome function as receptors for ubiquitin-modified proteins (Berko et al., 2014; Hofmann and Falquet, 2001; Husnjak et al., 2008; Mayor et al., 2005; Paraskevopoulos et al., 2014; Rosenzweig et al., 2012; Rubin et al., 1997; Saeki, 2017; Shi et al., 2016). In addition Rad23, Dsk2 and Ddi1 shuttle ubiquitin-modified proteins to the proteasome. A hallmark feature of such protein is that they carry a UBA domain that binds to ubiquitin chains on the substrates, and a UBL domain that binds to the proteasome (Chen and Madura, 2002; Kang et al., 2006). These domains architecture allows the shuttle factors to bridge the substrate and the proteasome efficiently. Till now such an ubiquitin binding domain has not been identified in Ccr4. Interestingly, we showed that the Ccr4 protein binds to 19S regulatory particle of the 26S proteasome (Figure 7) and (Laribee et al., 2007). We therefore asked if Ccr4 would bind to intracellular ubiquitin conjugates. Intriguingly, our IP experiment showed that Ccr4 indeed binds to cellular ubiquitin conjugates (Figure 5B) under native conditions. Upon de-naturation of ubiquitin conjugates, Ccr4 was unable to bind ubiquitin suggesting that Ccr4 requires native conformation of ubiquitin chains for its binding (Figure 6). Furthermore, the same result also excluded the possibility that Ccr4 itself being ubiquitin modified *in vivo*. Our next analysis revealed that LRR and nuclease domain of Ccr4 protein is essential for it ubiquitin shuttle function (Figure 8) and the exact residues and the regions of the LRR and nuclease domain of Ccr4 responsible for proteasome as well as ubiquitin chain binding remains to be explored and might be present within the LRR and C terminal part of Ccr4 protein.

**Figure 7:**
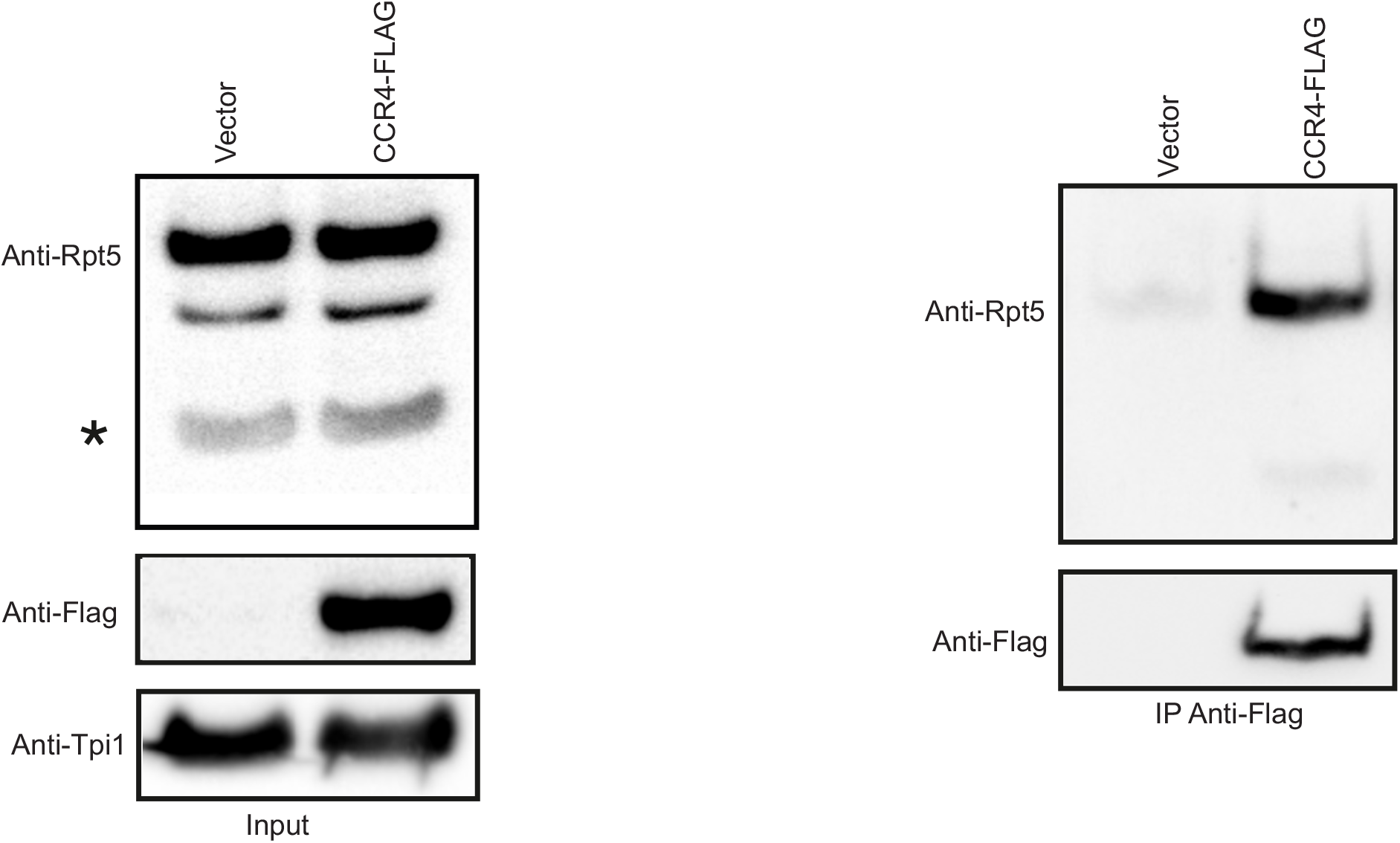
Ccr4 binds to 19S regulatory particle subunit complex. Western blot showing the association of Rpt5 subunit with 3X-FLAG-Ccr4. CO-IP experiment was performed as described in Figure 5B and also refer methods section. Input levels (left panel) and IP eluate (right panel) were analyzed by western blots using anti-Rpt5 and anti-Flag antibodies. Tpi1 was used as loading control. Asterisk denotes degradation product of Rpt5.

**Figure 8:**
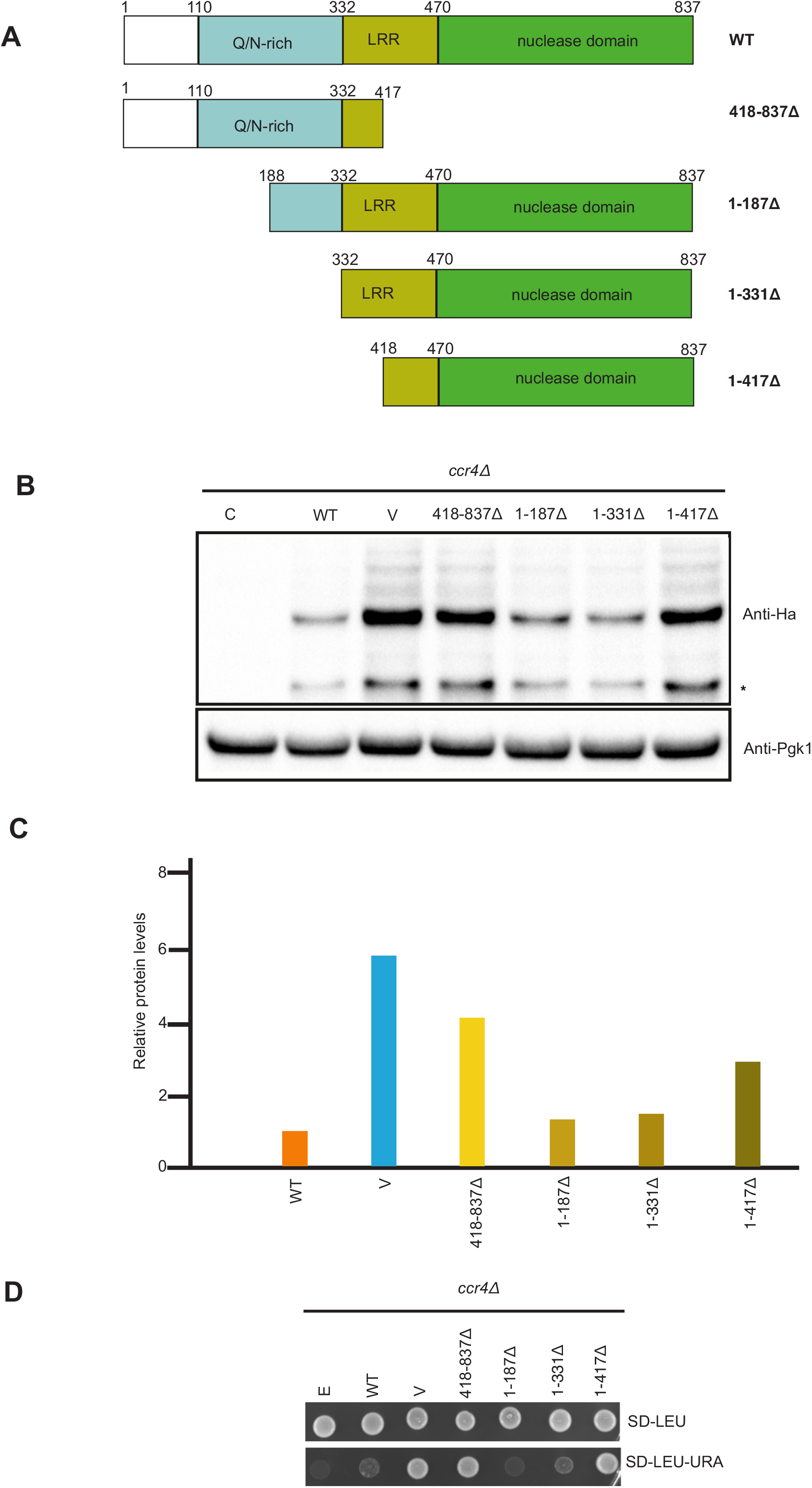
Effects of CCR4 truncations on UFD substrate degradation. (A) Cartoon illustrating different truncation of yeast Ccr4 protein. (B) Steady state levels of UFD substrate (Ub-V^76^-Ura3) levels in various CCR4 truncation mutants. (C) Densitometric analysis of Ub-V^76^-Ura3 protein levels. (D) Growth assay showing the accumulation of Ura3 based UFD substrate protein. *ccr4Δ* mutant strain carrying either empty plasmid vector or plasmid expressing either full length CCR4 or its different truncation variants were co-transformed with pBA2VV plasmid. Stabilization of UFD substrate was monitored based on the growth phenotype on Uracil lacking medium. cells were spotted on selection plates as indicated and photographed as described earlier in figure 3B. Anti-Ha and anti-Pgk1 antibodies were used for western blotting.

Our findings indicate that there are still unknown ubiquitin shuttle factors involved in targeting ubiquitin-modified substrates to the proteasome. The same conclusion was derived from a previous study in which yeast cells lacking multiple known shuttle factor shown to be viable (Husnjak et al., 2008). Our finding revels such a novel shuttle factor that has different structural characteristics than the known UBA-UBL domain containing proteins (Kang et al., 2006). Interestingly, Ccr4 protein is a highly conserved protein from lower to higher eukaryotes. Based on its structural and functional conservation we postulate that Ccr4 is involved in shuttling ubiquitin conjugates to the proteasome also in mammals. However such interesting hypothesis needs experimental validation and that is beyond the scope of this study.

Based on our understanding of the role of Ccr4 in protein degradation we have developed a model as shown in Figure 9. There is an intriguing list of shuttle factors for targeting of ubiquitin-modified substrates that are described so far. Interestingly, there is an additional layer of substrate selectivity at the substrate-targeting step to the proteasome. Almost all the shuttle factors have preference to bind a particular ubiquitin chain type having a suitable conformation for efficient recognition by the UBA domain or other ubiquitin receptors at the proteasome (Kang et al., 2006). Interestingly, Ccr4 may have preference to certain ubiquitin chain types for targeting. Therefore, it might have role beyond the UFD pathway to target other substrates of the proteasome for degradation. Based upon our findings such interesting possibilities can be explored for further understanding of Ccr4 functions in the UPS. Importantly, many diseases are associated and/or are caused by defects in the protein degradation our enhanced understanding of targeting by Ccr4 with in the UPS may help in finding a therapeutic approaches for such diseases in the future.

**Figure 9:**
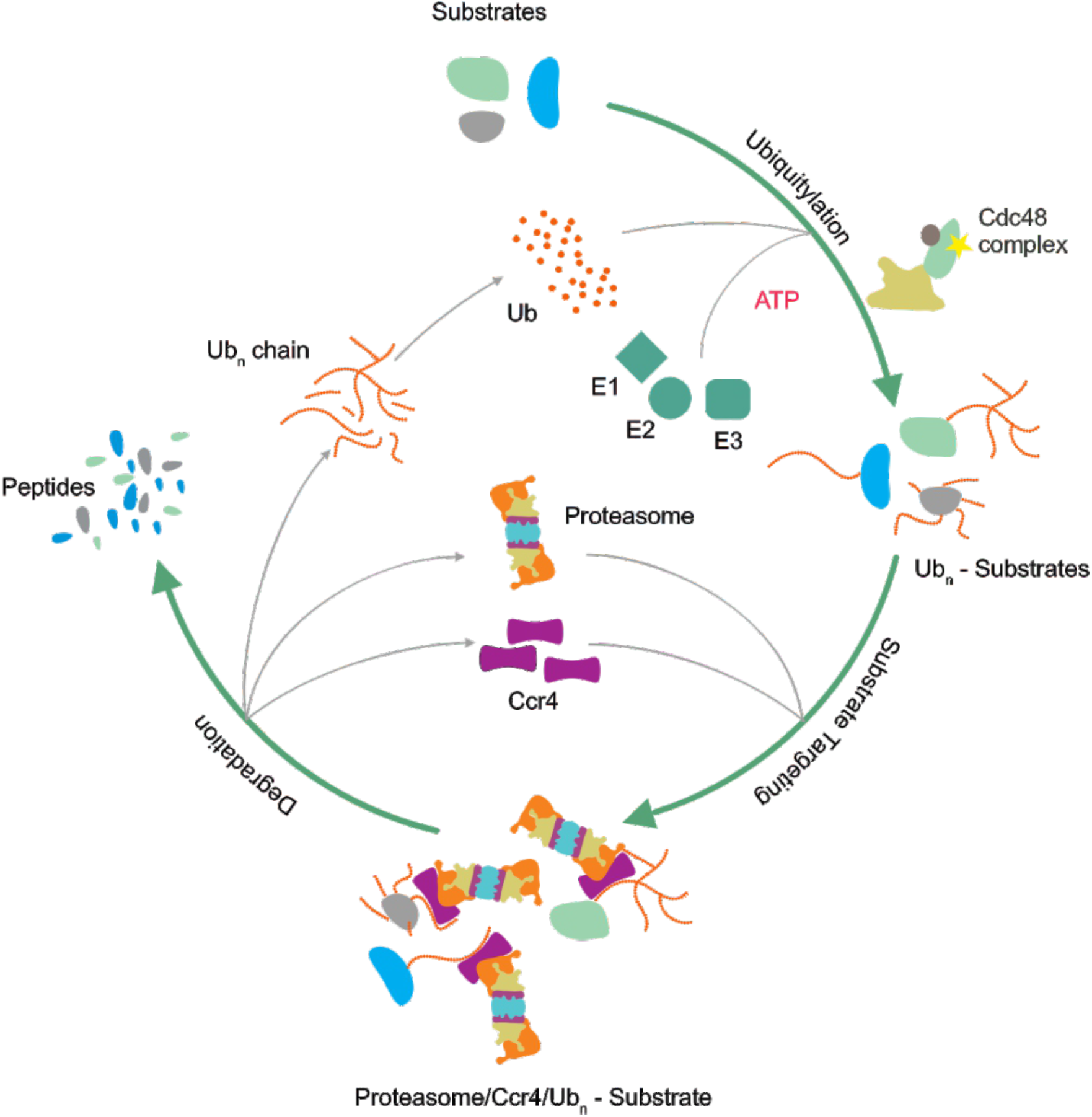
Ccr4 is an ubiquitin-modified protein targeting factor. Model showing the UFD pathway in eukaryotes. UFD substrate is ubiquitin modified by Uba1(E1), Ubc4/5(E2s), Ufd4(E3) then Cdc48^Npl4-Ufd1-Ufd2^ optimizes the ubiquitin chain. At a post-ubiquitylation step Ccr4 binds on the one hand binds to the proteasome and on the other hand Ccr4 binds to the ubiquitin conjugates thereby promotes targeting of ubiquitin-modified proteins to the 26S proteasome for degradation.

## Acknowledgment

This work was supported by funding from DBT-Ramalingaswami fellowship contingency grant as well as extra mural grant from SERB-DST to PMR. We thank Jürgen Dohmen, Marcel Fröhlich and Kerstin Nürrenberg (Institute for Genetics, University of Cologne, Germany), Thomas Langer (Max-Planck Institute for Biology of Ageing, Cologne, Germany), Ana-Mafalda Escobar-Henriques Dias and Ramona Schuster (Institute for Genetics, University of Cologne, Germany), Claes Andréasson (The Wenner-Gren Institute, Stockholm University, Sweden), Krishnaveni Mishra (University of Hyderabad, India) for strains as well as plasmids. We also thank Sanjay Suman (CSIR-Center for Cellular and Molecular Biology, Hyderabad, India) for help during this work. We thank Revathi Muthuvel for generating the polyclonal anti-Tpi1 antibody used in this study and Aswani Kumar for his help in preparing the illustrations.

## Supplemental Information

**Supplementary Figure 1:**
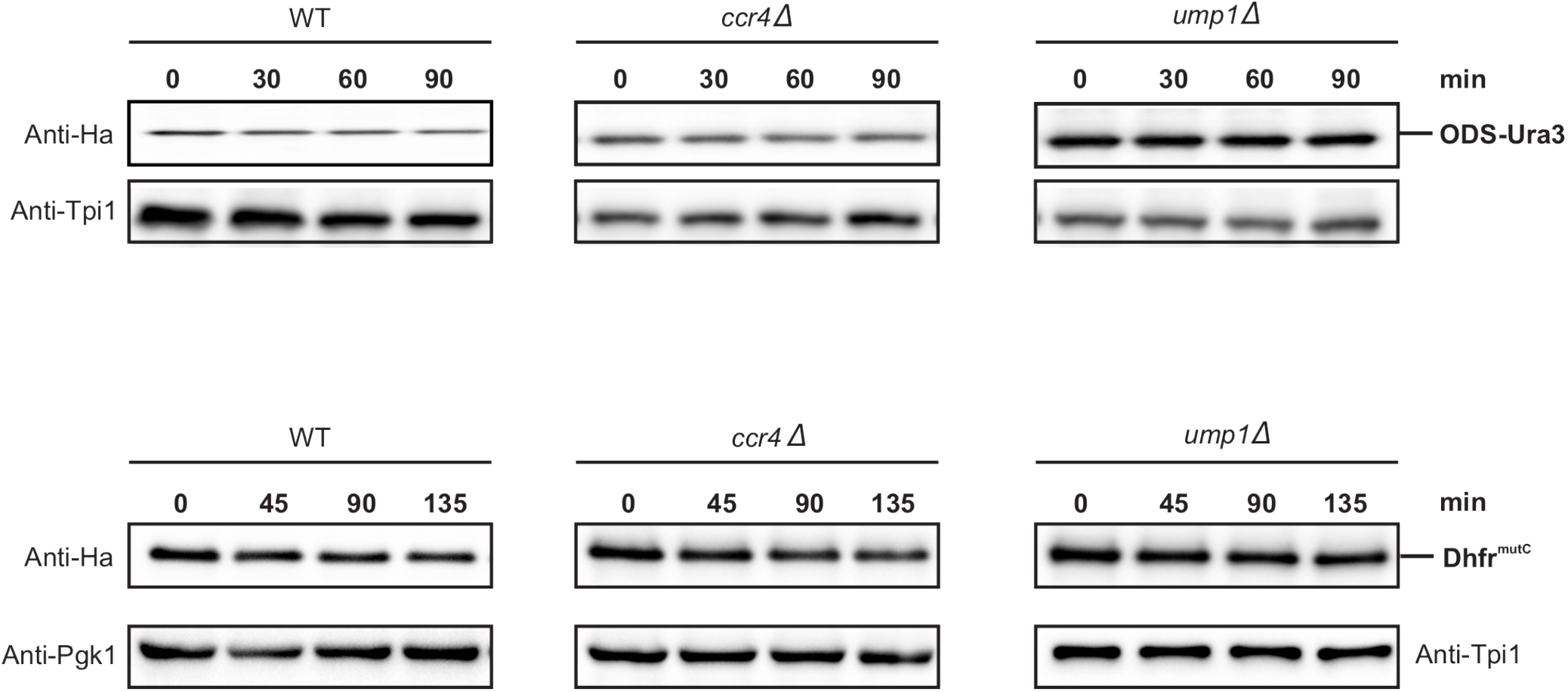
Cycloheximide chase assay for ODS-Ura3 and Dhfr^mutC^ substrate degradation. (A). WT, *ccr4Δ* and *ump1Δ* strains were transformed with pDG258 and pMAF56C. The cells were grown in SD-leu with 100 μM CuSO_4_ till mid-log phase at 25°C. In the next step cycloheximide chase was done as mentioned in methods section and the samples were analyzed by western blotting. Anti-Ha was used for detecting ODS-Ura3 and Dhfr^mutC^ (upper panels). Tpi1 and Pgk1 served as loading controls (lower panels).

**Supplementary Figure 2:**
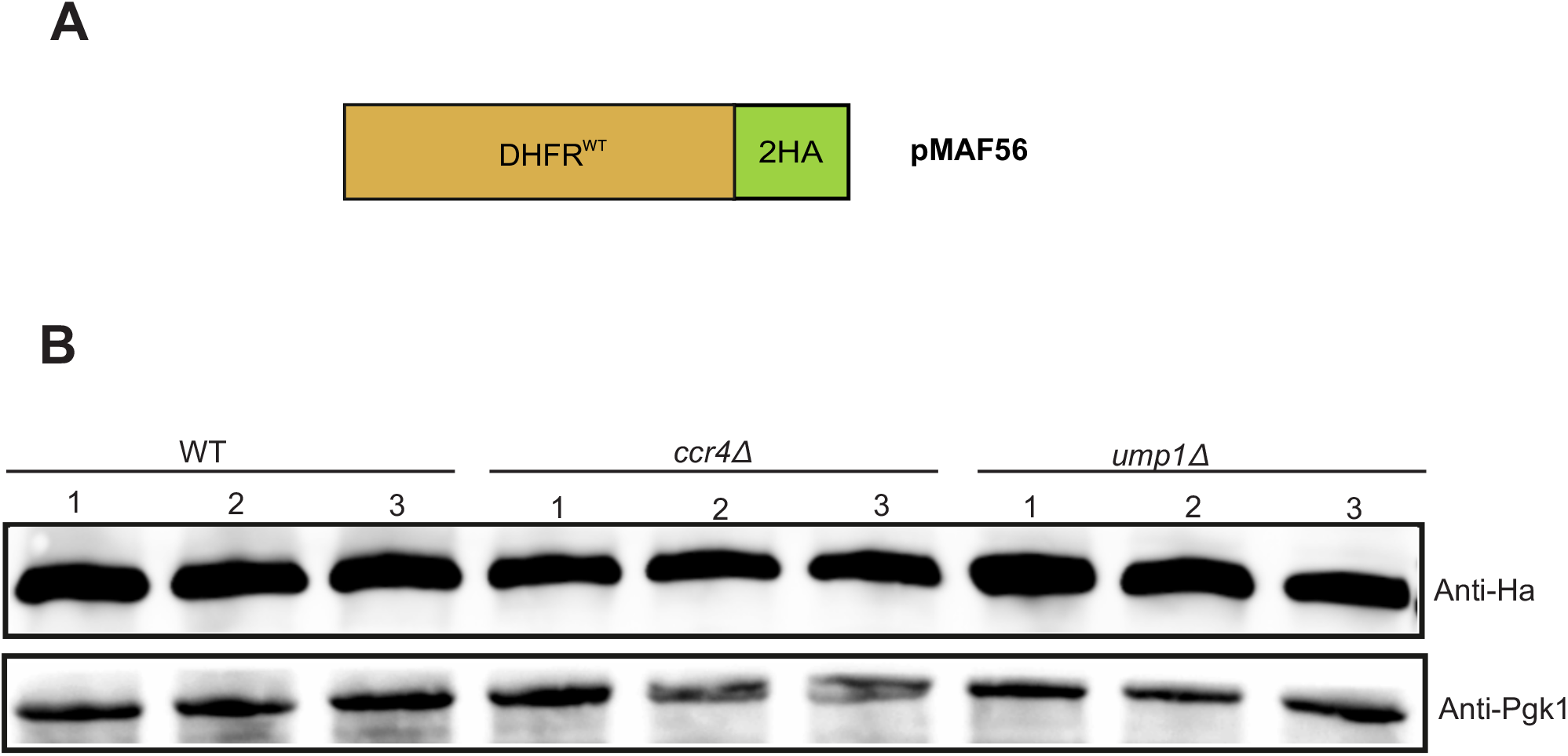
Steady state levels of Dhfr^WT^ substrate. (A). Schematic represents construct details of pMAF56 plasmid producing stable Dhfr wild-type protein. (B). Western blot showing the steady state levels of Dhfr^WT^ protein in WT, *ccr4Δ* and *ump1Δ* strains. Anti-Ha and anti-Pgk1 antibodies were used for immunoblot analysis. Data shown is the biological replicates of three independent samples.

